# Identification of glioblastoma stem cell-associated lncRNAs using single-cell RNA-sequencing datasets

**DOI:** 10.1101/2023.01.20.524887

**Authors:** Rasmani Hazra, Raditya Utama, Payal Naik, Alexander Dobin, David L. Spector

**Affiliations:** Cold Spring Harbor Laboratory, Cold Spring Harbor, NY 11724, USA

**Keywords:** lncRNA, glioblastoma, glioblastoma stem cell, single-cell RNA sequencing, radial glial

## Abstract

Glioblastoma multiforme (GBM) is an aggressive, heterogeneous grade IV brain tumor. Glioblastoma stem cells (GSCs) initiate the tumor and are known culprits of therapy resistance. Mounting evidence has demonstrated a regulatory role of long non-coding RNAs (lncRNAs) in various biological processes, including pluripotency, differentiation, and tumorigenesis. A few studies have suggested that aberrant expression of lncRNAs is associated with GSCs. However, a comprehensive single-cell analysis of the GSC-associated lncRNA transcriptome has not been carried out. Here, we analyzed recently published single-cell RNA-sequencing datasets of adult human GBM tumors, GBM organoids, GSC-enriched GBM tumors, and developing human brains to identify lncRNAs highly expressed in GBM. To categorize GSC populations in the GBM tumors, we used the GSC marker genes SOX2, PROM1, FUT4, and L1CAM. We found three major GSC population clusters: radial glia, oligodendrocyte progenitor cells, and neurons. We found 10–100 lncRNAs significantly enriched in different GSC populations. We also validated the level of expression and localization of several GSC-enriched lncRNAs using qRT-PCR, single-molecule RNA FISH, and sub-cellular fractionation. We found that the radial glia GSC-enriched lncRNA *PANTR1* is highly expressed in GSC lines and is localized to both the cytoplasmic and nuclear fractions. In contrast, the neuronal GSC-enriched lncRNAs *LINC01563* and *MALAT1* are highly enriched in the nuclear fraction of GSCs. Together, this study identified a panel of uncharacterized GSC-specific lncRNAs. These findings set the stage for future in-depth studies to examine their role in GBM pathology and their potential as biomarkers and/or therapeutic targets in GBM.

## Introduction

Glioblastoma multiforme (GBM, a World Health Organization grade IV glioma) ranks as the deadliest primary malignant brain cancer, with nearly 25,700 new cases diagnosed each year in the United States [1]. The currently available treatment regimen comprises maximal surgical resection, and a combination of radiotherapy and chemotherapy which extends patient survival to a median of only 14.6 months [2]. Despite recent progress in understanding the tumor’s biology and multimodal therapy options, GBM remains one of the most treatment-resistant malignancies, and even after successful treatment, the tumor inevitably recurs. Intratumor heterogeneity and therapeutic resistance in GBM are thought to be promoted by a cell population with stem cell-like properties, including self-renewal and differentiation [3–5]. GSCs are characteristically similar to normal neural stem cells, as they possess self-renewal capacity and express the CD133 gene (PROMININ-1), a marker for normal neural stem cells [5]. The radio-resistant and chemo-resistant GSC populations in GBM have been successfully isolated from GBM patient samples using cell surface membrane markers, including CD133 [5–7], CD44 [8], CD15 (SSEA1) [9], CD49f [10], L1CAM [11], PDGFRA [12], EGFR [13], and A2B5 [14]. Recent advances in technology, including single-cell RNA-sequencing (scRNA-seq) and lineage tracing techniques, have provided further evidence for the existence of GSC populations in GBM tumors [15]. An increasing body of evidence has shown that GSCs play an indispensable role in tumor initiation, progression, treatment resistance, and recurrence [5, 16, 17] indicating that GSCs are a critical therapeutic target.

Until relatively recently, proteins have traditionally been thought to be the critical regulators of biological processes; however, there is mounting evidence that non-coding RNAs also play important regulatory roles in almost all biological processes [18–20]. Advancements in sequencing technologies have enabled the discovery of large numbers of non-coding RNAs in the mammalian genome, which are transcribed in the cells but whose resulting transcripts do not code for proteins [21–23]. Long non-coding RNAs (lncRNAs) are defined as transcripts greater than 200 nucleotides in length with no protein-coding potential due to their lack of an open reading frame [24, 25]. They are transcribed by RNA polymerase II in the sense or antisense orientation and commonly originate from intergenic regions [26]. LncRNAs have been implicated in numerous regulatory molecular functions, including modulating transcriptional patterns, regulating protein activities, playing structural or organizational roles, altering RNA processing events, and serving as precursors to small RNAs [27–30]. Many lncRNAs are expressed in a tissue-specific manner, and some show significant changes in expression during differentiation [28, 31]. Moreover, altered lncRNA expression has been observed in several diseases, including cancer [32].

Individual lncRNA expression is often highly restricted to particular brain regions, and it has been suggested that lncRNAs offer more information about cell type identity during mammalian cortical development than protein-coding genes [33]. In addition, recent studies have shown that normal neurodevelopment in the human brain mirrors GBM development [34]. Therefore, the vast repertoire of lncRNAs in normal brain tissue indicates their potential importance in the context of GBM heterogeneity and treatment resistance. However, despite the large number of lncRNAs expressed in the brain, thus far just a handful have been shown to be involved in GSC biology. For example, the most abundant lncRNA, *MALAT1*, regulates proliferation and stemness in GSCs by activating the ERK/MAPK signaling pathway [35]. Knockdown of *XIST* in GSCs decreases proliferation, migration and invasion via upregulation of miR-152 [36]. Another abundant lncRNA, *NEAT1*, is highly expressed in GSCs. Depleting *NEAT1* inhibits GSC proliferation, migration, and invasion via upregulation of let-7e expression and downregulation of NRAS protein [37]. Down-regulation of *SOX2OT* in GSCs reduced their proliferation, migration, and invasion, and induced apoptosis in GSCs [38]. In addition, the maternally expressed lncRNA *H19* is highly expressed in GSCs and GBM tissue and is involved in maintaining GSC stemness and proliferation. However, a detailed functional analysis of this mechanism has not been elucidated [39]. Given these few proof-of-concept examples, a deeper screening and identification/characterization of novel lncRNAs is required to identify appropriate GSC-specific targets in GBM.

Here, we identified a large number of presently uncharacterized lncRNAs associated with GSC populations in GBM. We performed a systematic bioinformatics analysis using three scRNA-seq databases of GBM patient tumors, GBM patient-derived organoids and normal developing human brain [40–42]. We observed three main cell populations: radial glia, oligodendrocyte precursor cells (OPCs), and neuron populations. Furthermore, we identified GSC and non-GSC cell types in every cluster using the cancer stem cell markers CD133, CD44, CD15, FUT4, L1CAM, and SOX2 [3]. Using this analysis, we identified a large number of presently uncharacterized lncRNAs enriched with the GSC populations. Next, we validated the expression and subcellular localization of three selected lncRNAs using patient-derived GSC lines by qRT-PCR, single-molecule RNA fluorescence in situ hybridization (FISH), and subcellular fractionation. In summary, our analyses identified a large number of lncRNAs associated with the GSCs transcriptome and which should be pursued at the functional level and potentially as therapeutic targets in GBM.

## Methods

### Dataset preparation and sequencing for single-cell RNA-Seq

All datasets used in this study are publicly available. For single-cell RNA-Seq, the original main dataset [40] consisted of 32,877 cells from 11 GBM tumors obtained by surgical resection and processed immediately for RNA-seq. We also obtained a dataset [41] from glioblastoma derived organoids (GSE141947). For the validation of those previous datasets, we used a single cell RNA-seq dataset consisting of 3 GBM patient samples [43] (GSE139448). We further used a single cell RNA sequencing dataset [42] from developing normal human brain to compare lncRNA expression in normal vs GBM samples. We used RNA-seq glioblastoma patient-derived xenograft (PDX) GSC line datasets from the Mayo Clinic: (https://www.cbioportal.org/study/clinicalData?id=gbm_mayo_pdx_sarkaria_2019).

### Alignment and quantification for single-cell RNA-Seq

Sequence alignment and quantification was carried out using STAR v2.7.8a with STARsolo parameters. The primary single-cell GBM dataset [40], consisting of six aligned samples in BAM format (PRJNA579593), was processed with parameters as follows: (a) “--soloInputSAMattrBarcodeSeq CR UR” to specify cell and UMI sequences, (b) “-- soloInputSAMattrBarcodeQual CY UY” to specify cell and UMI quality and (c) “-- soloCBwhitelist 737-august-2016.txt” for 10X chromium v2. The single-cell organoid GBM dataset [41], consisting of 12 paired samples in FASTQ format (GSE141946), was processed with Drop-Seq parameters as follows: (a) “--soloCBwhitelist None” for Drop-Seq setting, (b) “-- soloCellFilter EmptyDrops_CR” to implement Cellranger v3 and v4 cell filtering method, (c) “-- soloCBstart 1” to specify cell barcode start position of 1, (d) “--soloCBlen 12” to specify cell barcode length of 12, (e) “--soloUMIstart 13” to specify UMI start position of 13, (f) “-- soloUMIlen 8” to specify UMI length of 8, (g) “--soloBarcodeReadLength 1” to specify the sum of cell and UMI barcode lengths. The human genome reference (FASTA assembly and GTF annotation) is from GENCODE GRCh38.p13.

### Pre-processing analysis for scRNA-Seq

The downstream analysis on scRNA-Seq data was mainly run on Scanpy v1.8.1. Doublet removal was done by Scrublet v0.2.3 to estimate number of droplets that isolate more than 1 cell (1205 cells removed). Data were filtered such that there was a minimum of three cells per gene (24797 genes removed) for minimum statistical calculation. To minimize low quality cell isolation and background contamination, we selected cells with a number of genes between 200 and 5,000 (631 cells removed). The maximum mitochondrial counts were set to 15% per cell for removing most dying cells. Log-normalized counts were calculated with a scale factor of 10,000 to generate gene expression values. We identified the top 2,000 highly variable genes with the “vst” method in Seurat v3 fashion, minimum mean expression of 0.0125, maximum mean expression of 3, minimum dispersion 0.5. Factors including total counts and mitochondrial percentage were regressed out from the expression data. Expression values were then scaled with maximum value of 10.

### Clustering analysis for scRNA-Seq

Local cell neighbors were calculated using 10 nearest neighbors (KNN) with the UMAP method, Euclidean distance, and 40 principal components. The Leiden clustering method, with a resolution of 2 (for the single-cell GBM dataset [40]) and 0.1 (for the single-cell organoid dataset [41]), was applied to optimize the coarseness of the clustering. Due to prominent sample-to-sample variation, samples were integrated with Harmony v0.0.5 to generate clusters associated with cell types.

### Dimensionality reduction for scRNA-Seq

Dimensionality reduction was performed using Principal Component Analysis (PCA) with the ARPACK method. Further, Uniform Manifold Approximation and Projection (UMAP) was calculated with a minimum distance of 0.2 (for the single-cell GBM dataset [40]) and 0.05 (for the single-cell organoid dataset [41]) to optimize the density of embedded cell points. Number of embedding dimensions was set to 2. Learning rate (alpha) and negative samples weight (gamma) were set to 1.

### Differential analysis for scRNA-Seq

Differential analysis was done using a Wilcoxon signed-rank test. Benjamini-Hochberg correction was applied to control for False Discovery Rate (FDR) and filter FDR<0.05.

### Alignment and quantification for bulk RNA-Seq

Sequence alignment was performed using STAR v2.7.8a with default parameters. The human genome reference (FASTA assembly and GTF annotation) was from GENCODE GRCh38.p13. Gene quantification was done by using feature Counts from Subread v2.0.2.

### Functional pathway analysis

Pathway analysis was accomplished by implementing GSEA v4.2.2 software. We collected pathways from widely used and curated databases, MSigDB v7.5.1including the Database for Annotation, Visualization, and Integrated Discovery (DAVID) gene ontology (GO) (for biological processes, cellular composition, and molecular function), Kyoto Encyclopedia of Genes and Genomes (KEGG), Reactome, and Hallmark. The whole gene set was pre-ranked and ordered by statistical scores, and the number of gene permutations was set to 1,000. The gene symbol remap was adapted from MSigDB v7.5.1.

### GSC lines

Patient-derived xenograft GSC lines (GSC #06, GSC #14, GSC #120, and GSC #161) were obtained from the Mayo Clinic [44] and cultured on laminin-coated plates (1–2 μg/cm^2^) (Sigma #L2020) in StemPro neural stem cell (NSC) medium. SFM is a serum-free medium (SFM) kit (Thermo Fisher Scientific #A1050901). Maintenance of the stem cell phenotype was verified by the expression of the stem cell markers SOX2, NESTIN, and CD133.

### Cell fractionation

Cell fractionation was performed as previously described [45]. Briefly, 3 million cells were resuspended in ice-cold lysis buffer (10 mM Tris pH 7.5, 150 mM NaCl, and 0.15% NP-40 substitute) for 15 minutes to lyse the cells. The cytoplasmic fraction was separated from the nuclei by overlaying the cell suspension on a sucrose buffer (10 mM Tris pH7.4, 150 mM NaCl, and 24% sucrose) and was centrifuged at 3,500 x g for 10 minutes. The remaining nuclear pellet was washed with ice-cold PBS-ethylenediaminetetraacetic acid (EDTA) and resuspended in urea buffer (1 M urea, 0.3 M NaCl, 7.5 mM MgCl_2_, 0.2 mM EDTA, and 1% NP-40) on ice for two minutes. The lysate was centrifuged at 13,000 x g for two minutes to separate the chromatin pellet from the nucleoplasm fraction supernatant. RNA was extracted from the cytoplasmic and nucleoplasmic fractions using TRIzol reagent (Sigma-Aldrich), according to the manufacturer’s protocol.

### RNA isolation and quantitative real-time PCR (qRT-PCR) assays

Total RNA was extracted from the cytoplasmic and nucleoplasmic fractions using TRIzol reagent (Sigma-Aldrich), according to the manufacturer’s instructions. Next, 1 μg of total RNA was used to synthesize cDNA using the TaqMan Reverse Transcription Reagent kit (Thermo Fischer). Then, 30 ng of cDNA was used to perform the qRT-PCR reaction using SYBR green PCR master mix on an ABI Quant Studio 6 Flex Real-Time PCR System (Applied Biosystems). The housekeeping genes GAPDH, PPIB, and PABPC1 were used as internal controls to normalize the gene of interest. Each experiment had two samples, and each assay was run three times. Primer sequences are listed below:

*GAPDH* (F: CACATCGCTCAGACACCATG and R: TCCCGTTCTCAGCCATGTAG)

*PPIB* (F: CAAGACAGACAGCCGGGATA and R: CTGTGGAATGTGAGGGGAGT)

*PABPC1* (F: GTTCGCAATCCTCAGCAACA and R: AAGAGTAGGGTGCATGGCTT)

*MALAT1* (F: CCCCTGGGCTTCTCTTAACA and R: TAGATCAAAAGGCACGGGGT)

*PANTR1* (F: CCCCAACCCAGTCACCTTAT and R: TTCCTGTGTTCACGTTGCTC)

*LINC01563* (F: TGAATACGGTGACTGAGGGG and R: AGGGCAGGAGAGAGGAAAAC)

### Single-molecule RNA fluorescence in situ hybridization (FISH)

Single-molecule RNA FISH was performed according to the manufacturer’s protocol for the RNAscope® Fluorescent Multiplex Reagent Kit 320850 (ACD #320850), as described previously [45, 46]. Briefly, 5 × 10^4^ GSCs were seeded onto laminin-coated coverslips (12 mm) in a 24-well plate for 24 hours to reach 80% confluence, then fixed in freshly prepared 4% paraformaldehyde (PFA) (Electron Microscopy Sciences, 19200). Fixed cells were permeabilized with different concentrations (50%, 80%, and 100%) of ethanol for 2 min, and 0.05% Triton X-100 for 10 min on ice before hybridization. The hybridization and signal amplification steps were performed according to the manufacturer’s instructions, and nuclei were counterstained with 4′,6-diamidino-2-phenylindole (DAPI). GSCs were imaged using a Zeiss LSM780 point scanning confocal microscope.

### Immunofluorescence

Immunofluorescence was performed using previously described protocols [45]. Briefly, GSCs were cultured on laminin-coated 12 mm glass coverslips for 24 hours to reach 80–90% confluence. The following day, the cells were fixed in freshly prepared 4% PFA for 30 minutes at room temperature, washed three times with phosphate-buffered saline (PBS), and permeabilized in PBS with 0.2% Triton-X-100 and 0.1% Tween-20 for 30 minutes on ice. Permeabilized cells were blocked in PBS with 2% horse serum for one hour at room temperature. Then the cells were incubated with anti-NESTIN (# ab22035, Abcam), anti-SOX2 (#ab 97959, Abcam) and anti-CD133 (#ab19898, Abcam) antibodies (dilution 1:100) overnight at 4°C. The next day, coverslips were washed three times in a blocking reagent and labeled with an anti-rabbit secondary antibody (Alexa fluor 488). After washing three times, the nuclei were counterstained with DAPI, and the coverslips were imaged using a Zeiss LSM780 point scanning confocal microscope.

### Overall Survival analysis

An online web tool Gene Expression Profiling Interactive Analysis (GEPIA2) (http://gepia2.cancer-pku.cn/), was used to predict overall survival analysis of selected lncRNAs in both low-grade glioma (LGG) and glioblastoma (GBM), This web server uses a comparative analysis using RNA-seq expression data from the TCGA. GEPIA uses the log-rank test (Mantel-Cox test), the Cox proportional hazard ratio, and a 95% confidence interval. The thresholds for high/low expression level cohorts can be adjusted.

## Results

### Identification of lncRNAs in multiple GSC populations in GBM tumors

To identify novel lncRNA transcripts associated with GSCs in GBM tumors, we analyzed a publicly available scRNA-seq dataset of GBM tumors [40]. This dataset sequenced 32,877 cells from 11 GBM tumors obtained by surgical resection and processed immediately for RNA-seq [40]. Due to stark differences in cell number, we selected cell clusters to maintain proper balance in the dataset and to avoid sample bias in the downstream analysis (Figure 1A). We ran sequence alignment, clustering, and visualization pipelines to generate optimal lists of differentially expressed lncRNAs for various cell types (detailed in Methods). UMAP-based dimensionality reduction improved cluster visibility compared to t-SNE which helped decompose cell types and states. To identify the known cell types for each GBM cluster, we further applied cell type markers (Figure S1A) provided by single nuclei transcriptomic analysis of developing human brain tissue [42]. From this analysis, we identified visually distinct main cell types enriched in given markers, including radial glia (identified by GFAP, HOPX), oligodendrocyte precursor cells (OPCs) (PDGFRA), neuronal (DLX1, DCX, NHLH1), microglia (C1QA), oligodendrocytes (MBP), astrocytes (GFAP), dividing cells (MKI67), pericytes (RGS5), endothelial cells (PECAM1), B cells (CD19), and red blood cells (GYPA) (Figure 1B and S1A). We identified a total of 1544 lncRNAs in GBM tumors. Of these 1544 lncRNAs, 549 lncRNAs are associated with radial glia, 328 lncRNAs are in the neuronal cluster, and 549 lncRNAs are associated with OPC clusters (Figure S1B and Table S1). Next, for every cell type, we classified cells as either GSC-positive [GSC (+)] or GSC-negative [GSC (-)] using the stemness markers PROM1, FUT4, or L1CAM, in conjunction with SOX2 and not expressing TLR4 [47] (Figure 1C). TLR4 is downregulated in GSCs [48] (Figure 1C). GSC-specific marker genes were extracted by implementing differential analysis on each cell type (see Methods). Given the low expression level of most lncRNAs, we filtered the list solely based on a significant false discovery rate (FDR), with values less than 0.05 (see Methods). Furthermore, our analysis observed a low number of defined GSCs due to the stringent combination of GSC markers. Therefore, we combined and thus reduced the number of clusters to increase statistical power in differential analysis. This analysis identified three GSC populations: radial glia, OPCs, and neurons. We identified 35 lncRNAs significantly enriched in the radial glia GSC population out of 391 differentially expressed lncRNAs. Among these lncRNAs, we found that uncharacterized lncRNAs including *LINC01088, ENSG00000288778, LINC02762*, and *gradually increased during hepatocarcinogenesis* (*GIHCG*) were highly expressed in the radial glia population (Figures 1D, 1E, S1C and Table S2). Similarly, 19 lncRNAs associated with GSCs were significantly enriched in the OPC population and 337 lncRNAs were enriched in the neuronal population (Figures 1D, 1E, S1C and Table S2). Interestingly, our clustering also revealed that the comparatively well-studied lncRNAs, such as *GAS5* is significantly enriched only in the radial glia GSC population but not in the OPC or neuronal GSC populations. On the otherhand, *NEAT1, NORAD, MEG8*, and *HOTAIRM1* lncRNAs are significantly enriched in both OPC and neuronal populations (Table S2). However, we observed very high expression *of MALAT1* in all three major cell types (radial glia, OPC and neuronal) (Table S1). We compared the lists of significant lncRNAs across cell types to generalize these findings. We observed only 6 overlapping lncRNAs among all three populations: radial glia, OPC, and neuron (Figures 1B and S1B). Further, we identified 16 common lncRNAs between the OPC and neuron populations, whereas 27 common lncRNAs were identified between the radial glia and neuronal populations, and 2 common lncRNAs were identified between the radial glia and OPC populations (Figure 1E and Table S2). We further *in-silico* validated this existing dataset with another publicly available dataset [43], where 12,367 cells were used from 3 patients. Using the same analyses, we have identified similar cell type populations, including three main clusters: radial glia, OPCs, and neuronal (Figure S1D).

**Figure 1:**
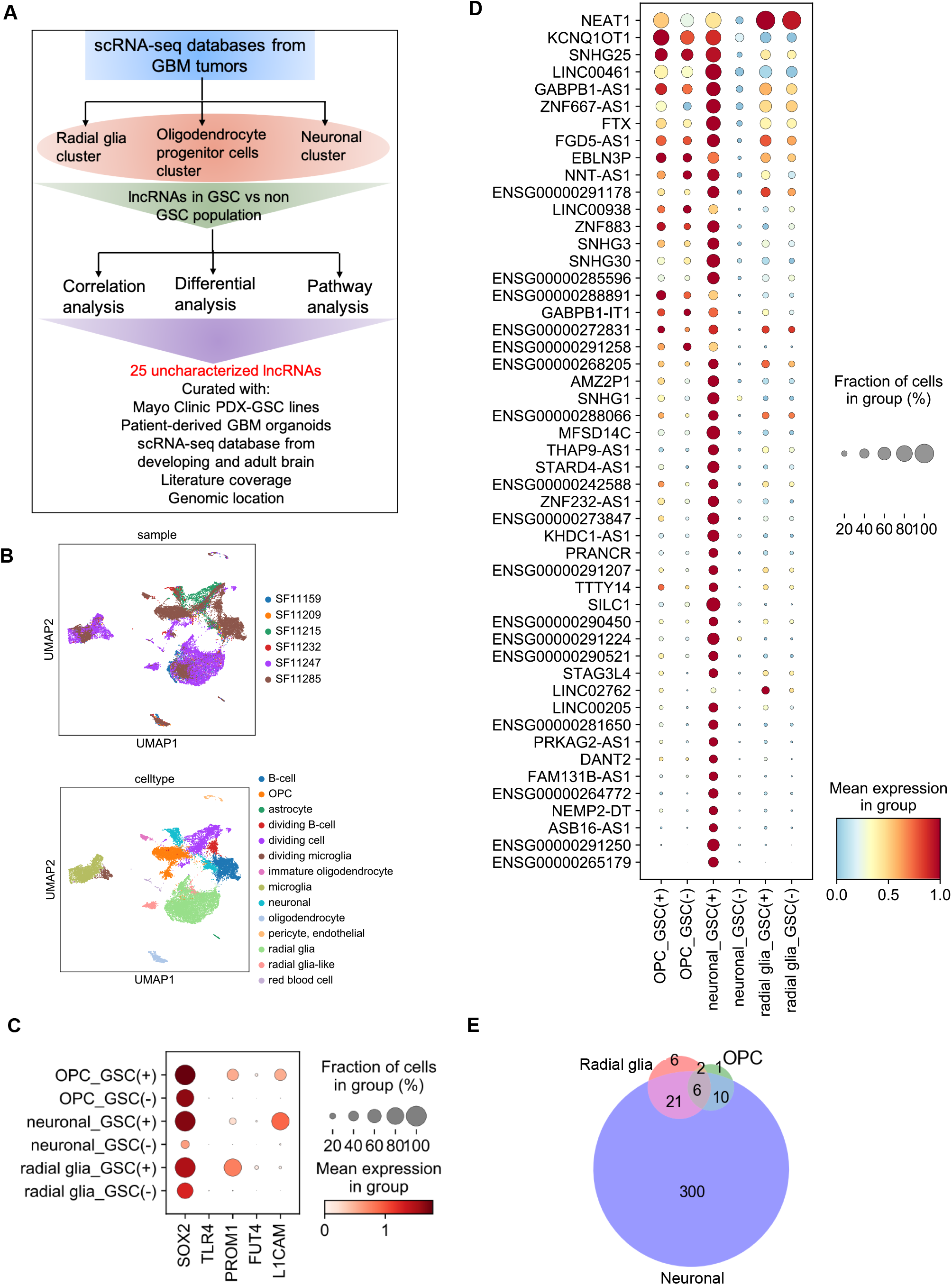
Identification of lncRNAs in multiple GSC populations in GBM tumors. (A) Workflow of GSC-enriched lncRNA analysis. Single-cell GBM dataset was reanalyzed to capture GSC populations in radial glia, oligodendrocyte precursor cells and neuronal clusters. Differential, pathway, and correlation analyses were performed on GSC(+) vs GSC(-) from which significant differential lncRNAs were identified. Quantitative and qualitative filtering were applied using various algorithms to identify enriched lncRNAs. (B) Samples composition and cell types were identified on UMAP using a combination of previously known markers including radial glia, oligodendrocyte precursor cells, neuronal cells, B-cells, dividing cells, endothelial, pericyte, oligodendrocyte, astrocyte, and red blood cells. (C) Expression and cellular fraction using GSC markers indicate a significant difference between GSC(+) and GSC(-) in the main 3 cell types. (D) List of 50 significant differentially expressed GSC lncRNAs in the 3 cell types summarized on a dot plot providing comparisons in scaled expression and cellular fraction. Genes were clustered by their scaled expression across cells. (E) Venn diagrams represent the number of overlapping and unique differentially expressed lncRNAs from all three cell populations.

### Comparative analyses of lncRNA expression levels across public GBM and GBO datasets

We next corroborated our uncharacterized GSCs-associated lncRNA set by performing a comparative analysis using scRNA-seq datasets from 8 GBM patient-derived organoid (GBO) lines, The Cancer Genome Atlas (TCGA), a bulk RNA-seq database of patient-derived xenograft GBM tumors, and a scRNA-seq dataset of the normal developing brain [42]. By implementing similar pipelines, we were able to identify similar cell-type clusters in the organoid dataset using identical cell-type markers, such as radial glia (GFAP, HOPX, and VIM), OPC (PDGFRA), and neuronal clusters (DCX and DLX1) (Figures 2A, 2B and S2A). We have also observed a highly distinct, dividing neuronal population identified by the MKI67 marker (Figure S2A). However, the expression of PDGFRA failed to capture a distinct OPC-like population (Figure S2A), which may be due to the selection in the cultured organoid medium. We next compared the sets of GSC (+) differentially expressed genes in radial glia, OPC, and neuron populations between GBM tumor and tumor-derived organoid datasets. Further, by comparing GBM tumor and organoid datasets, we found 27 common lncRNAs in the radial glia population (including *GIHCG, LINC01088, LINC027622*, and *SNHG6*) and only one common lncRNA (*MIAT*) in the neuronal GSCs and no common lncRNA in the OPC population (Figure 2B and Table S3). We observe a distinct radial glial population using the marker GFAP and HOPX in both tumor and tumor-derived organoid samples, however we did not observe distinct populations of OPC and neuronal GSCs in organoids using the markers PDGFRA (OPC) and DLX1, DCX, NHLH1 (neuronal). This limitation may be due to variation in selection in organoid culture.

**Figure 2:**
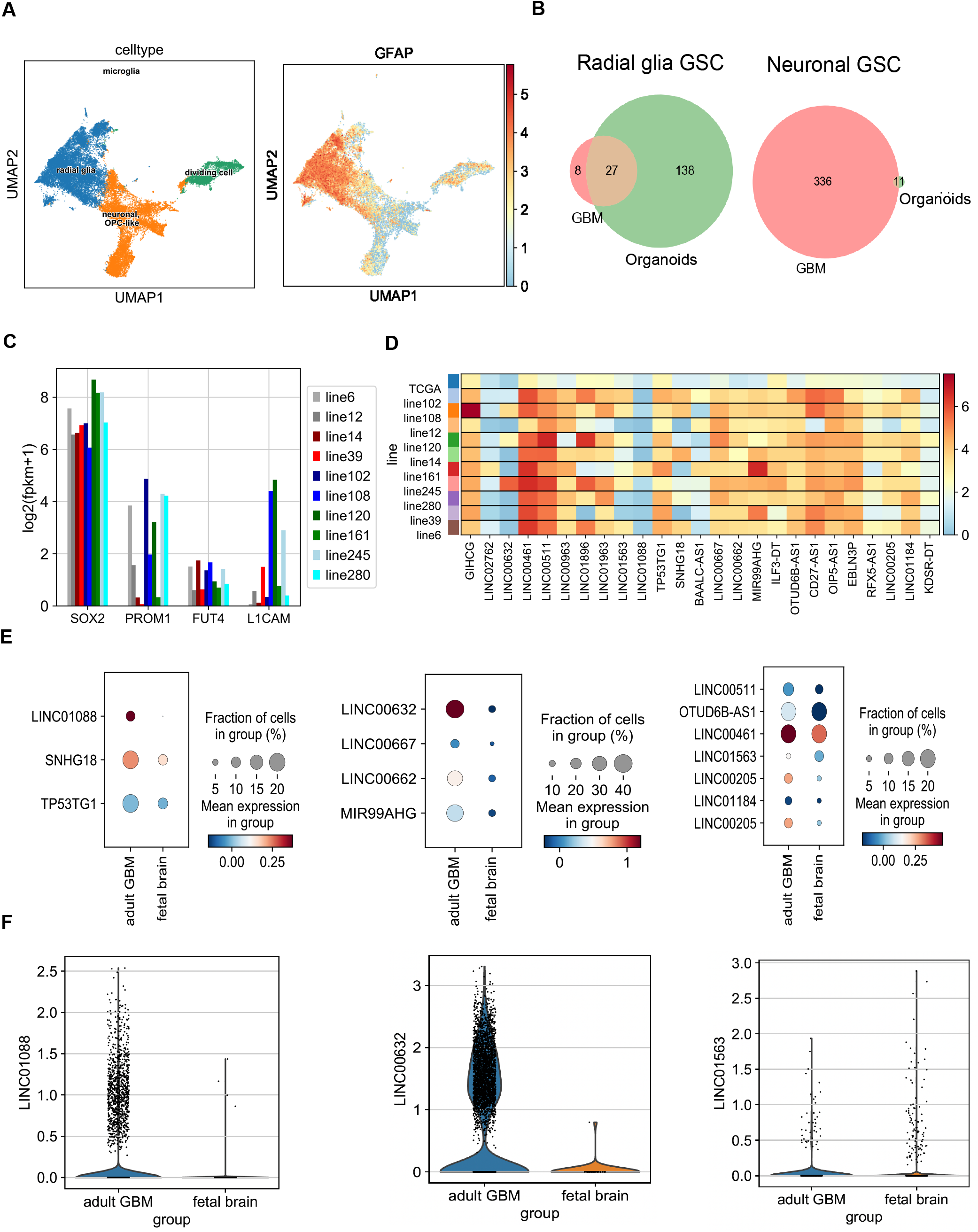
Comparative analyses of lncRNA expression levels across public GBM and GBO datasets. (A) Identification of a radial glial population in GBM-derived organoid dataset using known markers (GFAP). (B) Venn diagrams representing the number of overlapping and unique differentially expressed lncRNAs in GBM tissue and GBM organoids. (C) Expression levels of GSC markers were consistent across 10 Mayo Clinic cell lines from patient-derived xenograft (PDX) GBM samples. (D) Expression levels of 25 uncharacterized significant differentially expressed GSC lncRNAs are shown on heatmap across 10 Mayo Clinic cell lines and bulk TCGA GBM dataset. (E) Comparison between GBM and normal fetal brain are presented with the expression change and cellular fraction of differential GSC lncRNAs across 3 main cell types. (F) Expression of lncRNAs between GBM tumor and normal developing brain for radial glia, oligodendrocyte precursor cells and neuronal cell populations.

In general, the low expression of lncRNAs in most tissues, including GBM tissues, has been a great challenge in experimental validation. Thus, we selected the list of lncRNAs that have higher expression in tumor tissue and patient-derived xenograft (PDX) GSC lines using TCGA and patient-derived xenografts (PDX) bulk RNA-seq datasets [49]. We also showed that the GSC markers SOX2, PROM1, FUT4, and L1CAM were expressed in GSC lines (Figure 2C). Together with TCGA data, we selected the 25 uncharacterized intergenic lncRNAs that are differentially expressed in GSCs and have high expression levels (p < 0.05 and log2 (fold change) ≥ 2) (Figure 2D). Further, to investigate the expression levels of GSC-associated lncRNAs in the normal human brain, we compared the expression level between two independent datasets generated from adult GBM tumors [40, 43], and a normal developing human brain dataset [42]. It is known that developing human brain expresses thousands of lncRNAs and abnormal lncRNA expression has been associated with a wide range of neurological disorders [50]. Interestingly, we observed many of the lncRNAs identified from each of the three major clusters (radial glia, OPC, and neuronal) to be more highly expressed in the adult GBM tissue compared to the normal brain (Figures 2E, 2F, and S2C), suggesting that these lncRNAs are upregulated in the GSC populations in tumor but not in the normal human brain.

### Pathway analysis of GSC (+) cells suggests involvement of the radial glia population in metabolic-related processes

To further investigate the underlying biological functions of the GSC-enriched lncRNAs in GBM, Gene ontology (GO) term, Kyoto Encyclopedia of Genes and Genomes (KEGG), REACTOME, and HALLMARK pathway analyses were conducted using all genes ranked by statistical score (see Methods) generated from GSC vs non-GSC comparison separately in all three cell types (radial glia, OPC, and neuron). Interestingly, we did not observe any common pathways between all three GSC-associated cell types (Figures 3A, 3B, and 3C), suggesting the functional heterogeneity of GSC populations in GBM tumors. Biological pathways related to metabolic processes (oxidative phosphorylation, reactive oxygen species pathway, inner mitochondrial membrane protein complex, ATP synthesis coupled electron transport, NADH dehydrogenase activity) were significantly enriched in the radial glia GSC population, on the other hand, OPC and neuronal populations were enriched in different pathways involved in cancer development and progression, such as angiogenesis, epithelial-mesenchymal transition, apoptosis, MTORC1 signaling, hedgehog signaling, TGFB signaling, PI3K-AKT-MTOR signaling, G2M checkpoint, DNA repair, E2F targets (Figures 3A, 3B and 3C). Notably, the metabolic characteristics of GSC populations are highly heterogeneous in nature. The radial glia population is enriched in oxidative phosphorylation or mitochondrial metabolic pathway (Figure 3A). In contrast, the OPC population is enriched in the glycolytic pathway (Figure 3B), indicating the complex heterogeneous tumor microenvironment even within the same tumor.

**Figure 3:**
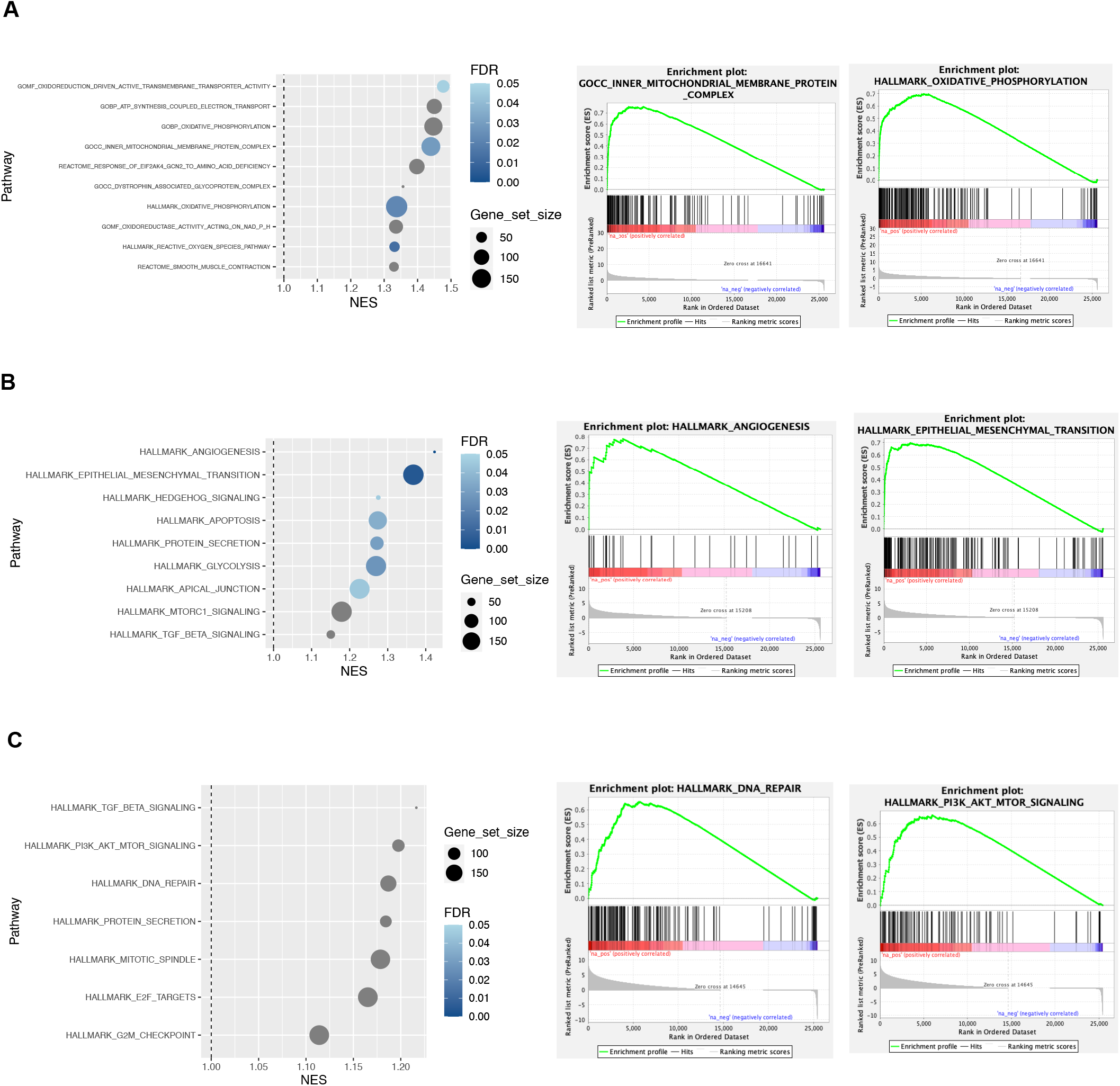
Pathway analysis of GSC (+) cells suggests involvement of the radial glia population in metabolic-related processes. (A) Radial glia GSC expression change is associated with pathways enriched in various metabolic functions including inner mitochondrial membrane protein complex and oxidative phosphorylation. (B) oligodendrocyte precursor GSC expression change is associated with pathways including angiogenesis and epithelial-mesenchymal transition (C) Neuronal GSC expression change is associated with pathways including TGF beta signaling, DNA repair, and PI3K-AKT-MTOR signaling.

### Validation of GSC-associated lncRNAs

We first validated the patient-derived xenograft GSC lines using established GSC markers (NESTIN, SOX2, CD133) [47]. Nestin is a class six intermediate filament proteins expressed in neural stem cells [51] and is expressed in the cytoplasm of GSC lines (Figures 4A and S4A). SOX2 is a transcription factor that plays a critical role in maintaining the self-renewal capability of neural stem cells. Its activity is associated with maintaining the undifferentiated state of cancer stem cells in several tissues [52]. We found that SOX2 is highly enriched in the nucleus of all four GSC lines (Figures 4A and S4A). CD133 (PROMININ1, PROM1) is a transmembrane glycoprotein commonly used as a marker of normal and cancerous stem cells, particularly in central nervous system tumors, including GBM [53]. On the other hand, CD133 (PROM1) is expressed in both the cytoplasm and the cell surface of GSC lines (Figures 4A and S4A). To validate our GSC-specific lncRNA expression patterns, we performed in situ hybridization for three lncRNAs: *MALAT1, LINC01563* and *PANTR1* (Figures 4B and S4B). Furthermore, we performed a cell fractionation assay followed by qRT-PCR, which showed that *PANTR1* was enriched in both the cytoplasm and nuclear fraction. In contrast, *LINC01563* and *MALAT1* were only enriched in the nuclear fraction (Figure 4C), further verifying the RNA-FISH signals in GSCs (Figure 4B). Survival analysis was also performed on these three lncRNAs. *LINC01563* significantly correlated with the overall survival of patients with both LGG and GBM (Figure 4D). *PANTR1* showed a survival prognosis for patients with LGG but not with GBM (Figures 4D and S4C). However, *MALAT1* lncRNA did not show a survival advantage in either LGG or GBM (Figure S4C).

**Figure 4:**
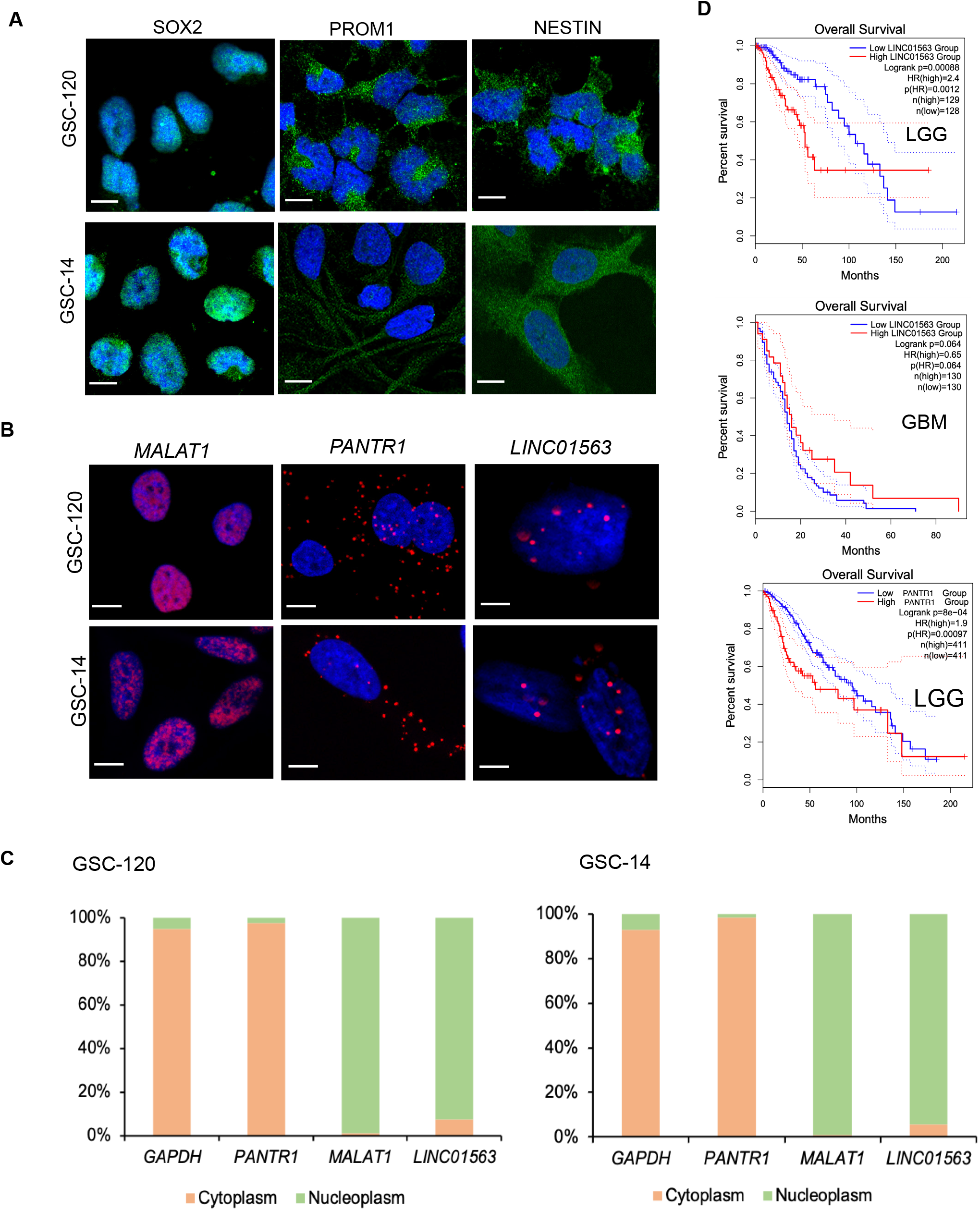
Validation of GSC-associated lncRNAs. (A) Immunolabeling of SOX2, PROM1 and NESTIN in GSC lines (GSC-120 and GSC-14). Scale bar, 12 μm. (B) Single-molecule RNA-FISH images indicate localization of *MALAT1, PANTR1*, and *LINC01563* lncRNA transcripts (red dots) within the nuclei and/or cytoplasm in GSCs. Scale bars, 12 μm. (C) Subcellular fractionation followed by qRT-PCR confirming the localization of *PANTR1* and *LINC01563* lncRNA transcripts. *GAPDH* and *MALAT1* were used as cytoplasmic and nuclear markers for quality control, respectively. Data are presented as mean values ± SEM (N=3 independent experiments). (D) Kaplan–Meier overall survival prognosis of *LINC01563* and *PANTR1*.

## Discussion

It is increasingly evident that lncRNAs act as biomarkers, tumor suppressors, and oncogenes in various cancers [54–57]. Recently, some in silico studies have identified lncRNAs in GBM tumors and cell lines using bulk RNA-seq from TCGA and microarray platforms [58–62], although very few of them have been characterized in-depth regarding their potential regulatory function in GBM. However, the bulk RNA-seq approach limits our understanding of GBM tumor heterogeneity, given its cellular complexity. Recent transcriptome analysis revealed that GBM tumors are highly heterogeneous [15]. Therefore, the scRNA-seq approach can identify different cell types in a single tumor. Since many lncRNAs are expressed in a cell-type specific manner, identifying novel lncRNAs in different cell-type clusters provides a means to study their regulatory role in GBM biology and may offer a down-stream lncRNA-based therapeutic approach.

Thus, we set out to provide a comprehensive analysis of the GSC transcriptome using scRNA-Seq databases of GBM tumors and tumor-derived organoids. While we confirmed several known cancer-associated lncRNAs such as *NORAD, NEAT1, MALAT1, XIST, PVT1, MEG3, MEG8* and *GAS5* in our screen, most of 1544 differentially expressed lncRNA candidates have not been described previously in the context of glioblastoma. In addition, we have identified for the first time 374 uncharacterized lncRNAs associated with GSCs. Our study emphasizes the importance of unbiased screening approaches and represents a valuable resource for GSC-associated lncRNAs to further study their regulatory role in GBM initiation, maintenance, and progression.

A DAVID GO term pathway analysis using GSC-pos and GSC-neg cells from three different clusters showed that differentially expressed genes were significantly enriched in different cancer-related pathways. Interestingly, the radial glia population was enriched in the electron transport chain and oxidative phosphorylation pathways and mitochondrial metabolism (Figures 4A and S4A), consistent with the recent findings that oxidative phosphorylation or mitochondrial metabolism are the preferred energy source for cancer stem cells (PMID: 34052208). On the other hand, the glycolytic pathway is significantly enriched in the OPC-GSC population, indicating that the metabolic state of GSCs differs in different cell populations. This finding demonstrates that metabolic characteristics in GSCs are highly heterogenous within the tumor. Studies have shown that cancer stem cells can switch their metabolic state to favor either oxidative metabolism or glycolysis, and this switch relies on the tumor microenvironment [63]. Thus, targeting GSCs metabolism by potentially manipulating lncRNA expression may provide new and effective methods for treating GBM tumors.

In conclusion, our in-silico comprehensive analysis provides a group of GSC-associated lncRNAs in GBM. Future in-depth mechanistic studies of these uncharacterized lncRNAs will extend the field’s understanding of GBM biology and offer new potential lncRNA biomarkers and therapeutic targets for GBM. Exploring the functional relevance of lncRNAs in GBM biology will provide better insights into this devastating disease and its potential treatment.

## Acknowledgments

We thank members of the Next-Gen Sequencing and Analysis Shared Resource for supporting the analysis. We also thank Andrea McMahon and Ann C.M. Tuma from Jann N. Sarkaria’s laboratory for providing us PDX-GSC lines. We also thank Tse-Luen (Erika) Wee, Director of the CSHL Microscopy Shared Resource for assistance with imaging.

## Author contributions

R.H. conceptualized and designed the research, performed the experiments, analyzed the data, and wrote the manuscript. R.U. did the bioinformatics analysis and wrote part of the Methods and Results sections of the manuscript, P.N. performed the experiments, A.D reviewed, and edited the manuscript, D.L.S. was responsible for acquisition of funding, reviewing, and editing the manuscript.

## Data Availability Statement

We did not generate any new datasets in this study, and we used publicly available datasets, which are mentioned in the Methods section.

## Figure Legends

**Figure S1:**
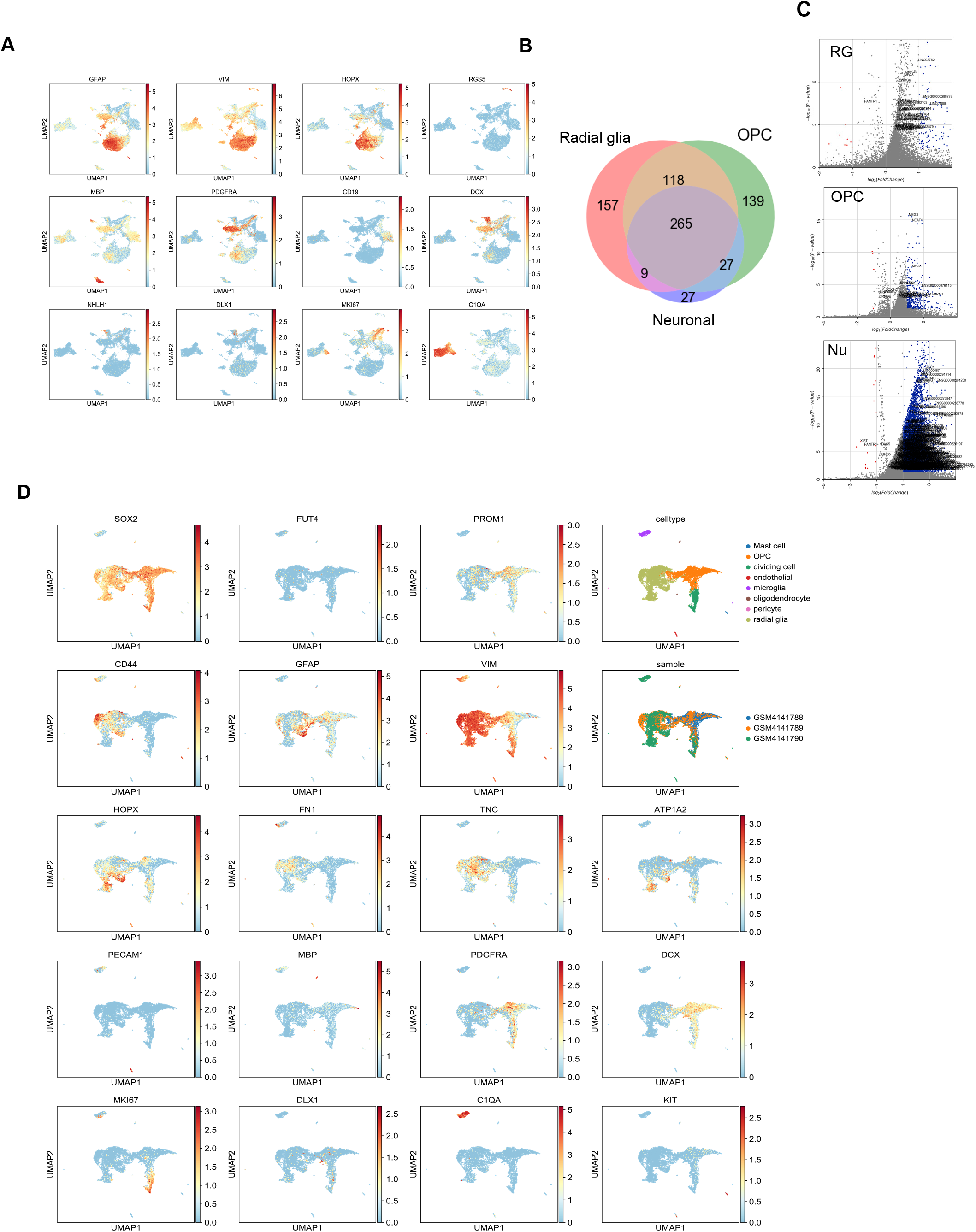
Identification of lncRNAs in multiple GSC populations in GBM tumors. (A) Sets of markers used to identify previously known cell types including radial glia (e.g GFAP, HOPX), neuronal (e.g DCX, DLX1), and OPC (e.g PDGFRA). (B) Differential GSC lncRNA statistics (p-value and logFC) in radial glia, oligodendrocyte precursor cells, and neuronal populations.

**Figure S2:**
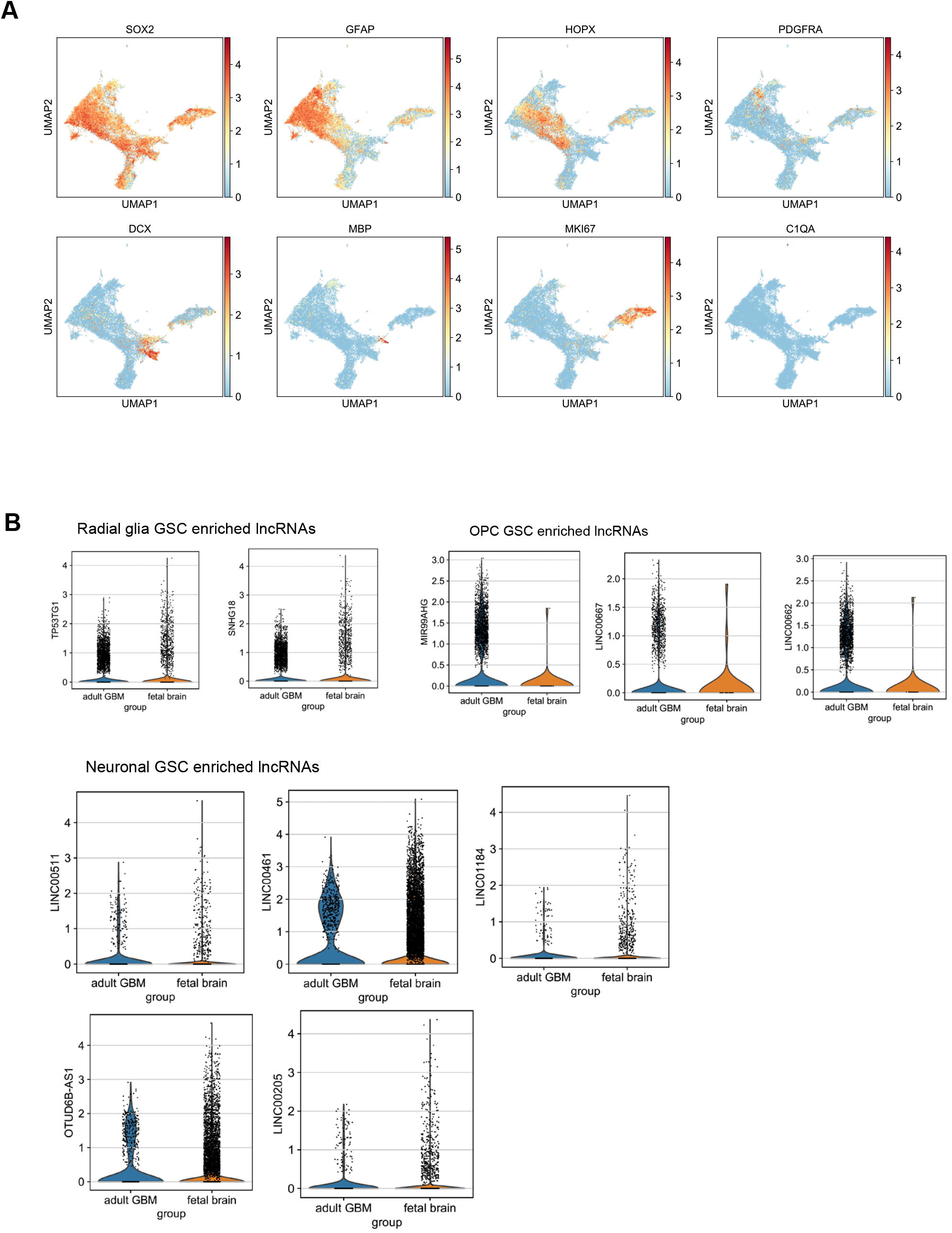
Comparative analyses of lncRNA expression levels across public GBM and GBO datasets. (B) Cell type markers used to identify previously known cell types including radial glia (GFAP, HOPX), neuronal (DCX, DLX1), and oligodendrocyte precursor cells (PDGFRA) in GBM-derived organoid dataset and (B) Expression of lncRNAs between GBM tumor and normal developing brain for radial glia, oligodendrocyte precursor cells and neuronal cell populations.

**Figure S3:**
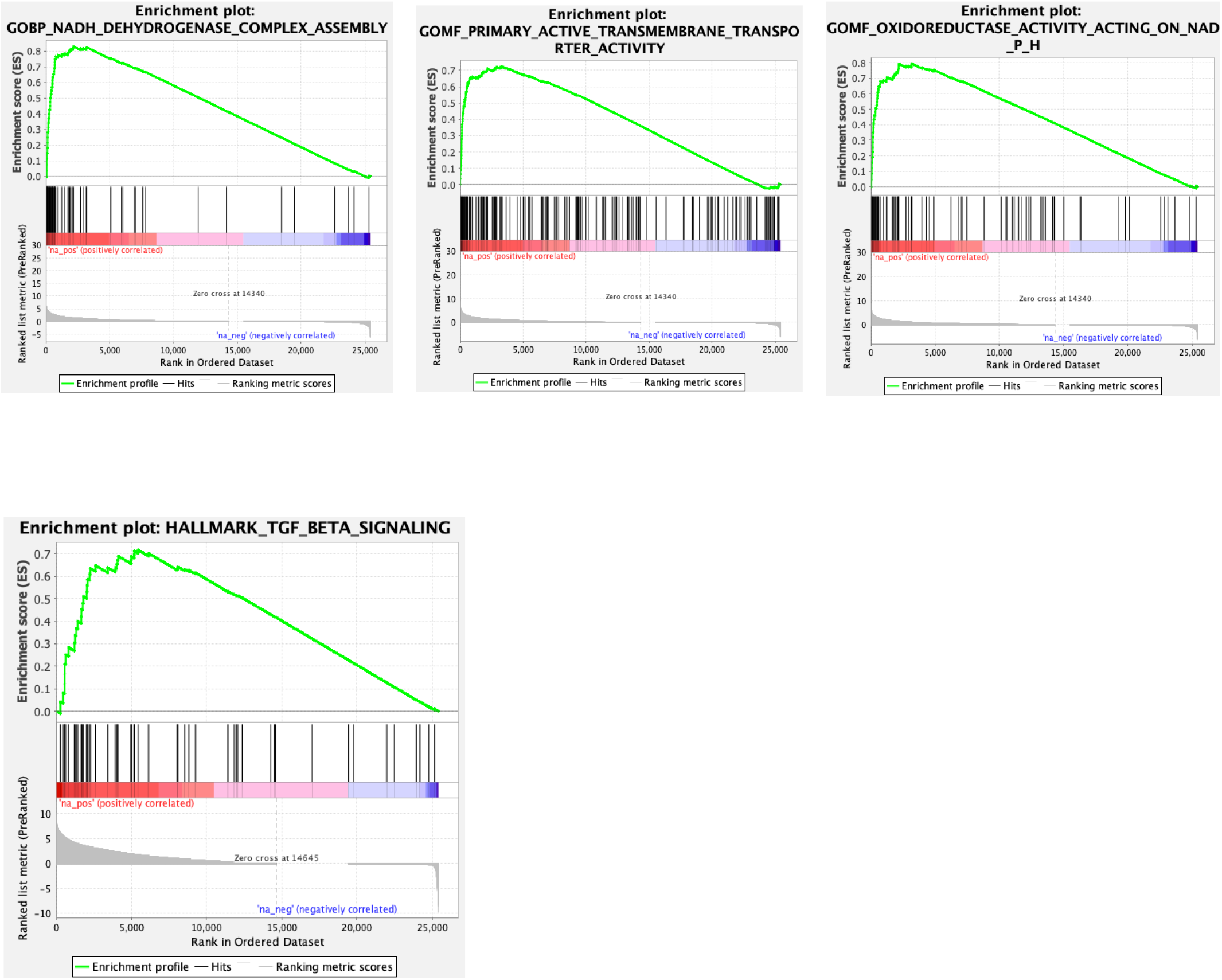
Pathway analysis of GSC (+) cells suggests involvement of the radial glia population in metabolic-related processes. GSEA pathway analysis showing pathways enriched in radial glial, OPC, and neuronal populations.

**Figure S4:**
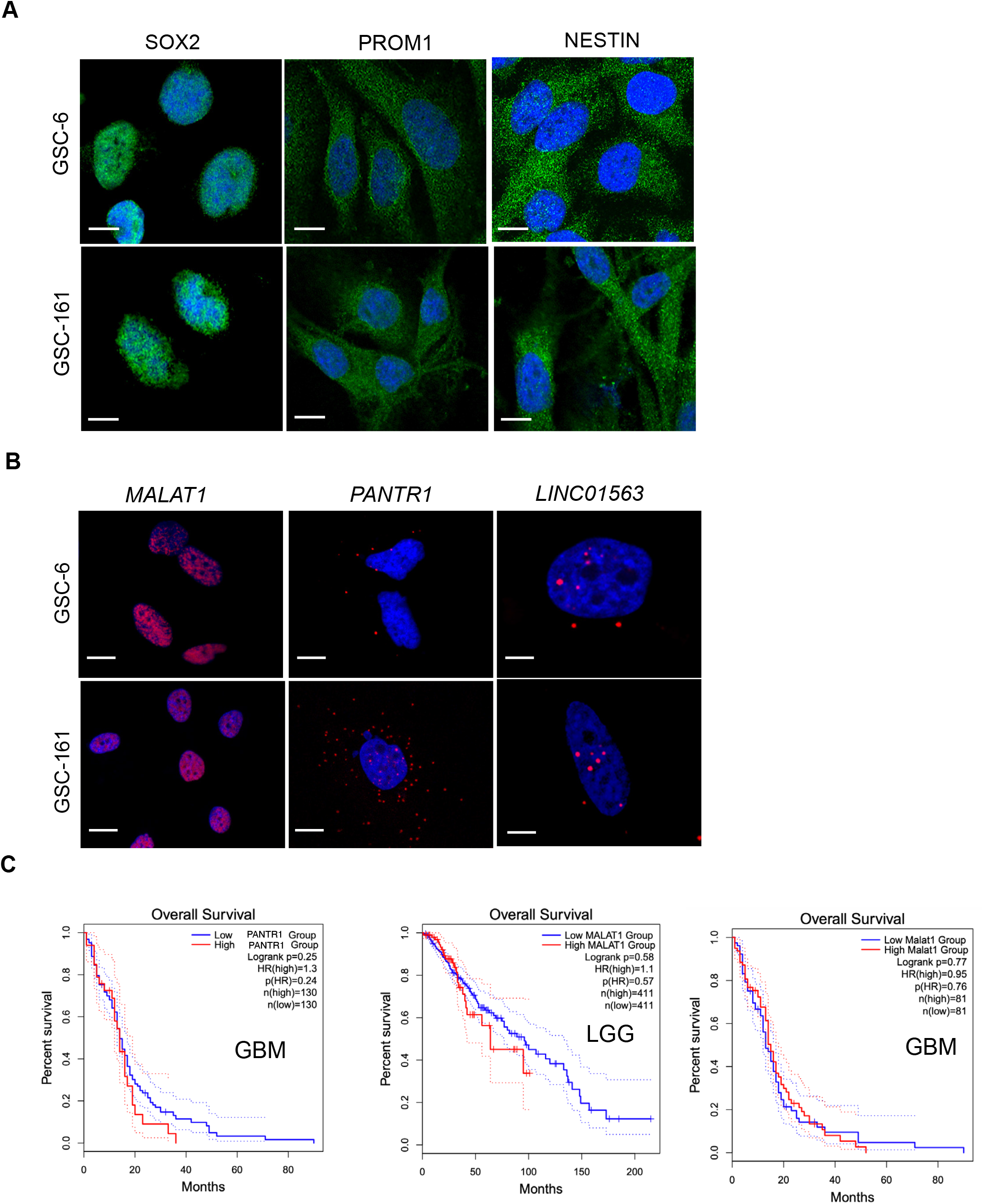
Validation of lncRNAs. (A) Immunolabeling of SOX2, NESTIN and CD133 in GSC lines (GSC-6 and GSC-161). Scale bar, 12 μm. (B) Single-molecule RNA-FISH images indicate localization of *MALAT1, PANTR1*, and *LINC01563* lncRNA transcripts (red dots) within nuclei and/or cytoplasm in GSC lines (GSC-6 and GSC-161). Scale bars, 12 μm. (C) Subcellular fractionation of GSCs followed by qRT-PCR confirmed the localization of *PANTR1* and *LINC01563* lncRNA transcripts. *GAPDH* and *MALAT1* were used as cytoplasmic and nuclear markers for quality control, respectively. Data are presented as mean values ± SEM (N=3 independent experiments). (D) No overall survival prognosis was identified for *MALAT1* and *PANTR1* in both LGG and GBM.

## References

1. Ostrom, Q.T., Cioffi, G., Waite, K., Kruchko, C., and Barnholtz-Sloan, J.S. (2021). CBTRUS Statistical Report: Primary Brain and Other Central Nervous System Tumors Diagnosed in the United States in 2014-2018. Neuro Oncol 23, iii1–iii105.

2. Prager, B.C., Bhargava, S., Mahadev, V., Hubert, C.G., and Rich, J.N. (2020). Glioblastoma Stem Cells: Driving Resilience through Chaos. Trends Cancer 6, 223–235.

3. Lathia, J.D., Mack, S.C., Mulkearns-Hubert, E.E., Valentim, C.L., and Rich, J.N. (2015). Cancer stem cells in glioblastoma. Genes Dev 29, 1203–1217.

4. Singh, S.K., Clarke, I.D., Terasaki, M., Bonn, V.E., Hawkins, C., Squire, J., and Dirks, P.B. (2003). Identification of a cancer stem cell in human brain tumors. Cancer Res 63, 5821–5828.

5. Singh, S.K., Hawkins, C., Clarke, I.D., Squire, J.A., Bayani, J., Hide, T., Henkelman, R.M., Cusimano, M.D., and Dirks, P.B. (2004). Identification of human brain tumour initiating cells. Nature 432, 396–401.

6. Lathia, J.D., Gallagher, J., Heddleston, J.M., Wang, J., Eyler, C.E., Macswords, J., Wu, Q., Vasanji, A., McLendon, R.E., Hjelmeland, A.B., et al. (2010). Integrin alpha 6 regulates glioblastoma stem cells. Cell Stem Cell 6, 421–432.

7. Piccirillo, S.G., Reynolds, B.A., Zanetti, N., Lamorte, G., Binda, E., Broggi, G., Brem, H., Olivi, A., Dimeco, F., and Vescovi, A.L. (2006). Bone morphogenetic proteins inhibit the tumorigenic potential of human brain tumour-initiating cells. Nature 444, 761–765.

8. Anido, J., Saez-Borderias, A., Gonzalez-Junca, A., Rodon, L., Folch, G., Carmona, M.A., Prieto-Sanchez, R.M., Barba, I., Martinez-Saez, E., Prudkin, L., et al. (2010). TGF-beta Receptor Inhibitors Target the CD44(high)/Id1(high) Glioma-Initiating Cell Population in Human Glioblastoma. Cancer Cell 18, 655–668.

9. Son, M.J., Woolard, K., Nam, D.H., Lee, J., and Fine, H.A. (2009). SSEA-1 is an enrichment marker for tumor-initiating cells in human glioblastoma. Cell Stem Cell 4, 440–452.

10. Barbar, L., Jain, T., Zimmer, M., Kruglikov, I., Sadick, J.S., Wang, M., Kalpana, K., Rose, I.V.L., Burstein, S.R., Rusielewicz, T., et al. (2020). CD49f Is a Novel Marker of Functional and Reactive Human iPSC-Derived Astrocytes. Neuron 107, 436–453 e412.

11. Bao, S., Wu, Q., Li, Z., Sathornsumetee, S., Wang, H., McLendon, R.E., Hjelmeland, A.B., and Rich, J.N. (2008). Targeting cancer stem cells through L1CAM suppresses glioma growth. Cancer Res 68, 6043–6048.

12. Kim, Y., Kim, E., Wu, Q., Guryanova, O., Hitomi, M., Lathia, J.D., Serwanski, D., Sloan, A.E., Weil, R.J., Lee, J., et al. (2012). Platelet-derived growth factor receptors differentially inform intertumoral and intratumoral heterogeneity. Genes Dev 26, 1247–1262.

13. Emlet, D.R., Gupta, P., Holgado-Madruga, M., Del Vecchio, C.A., Mitra, S.S., Han, S.Y., Li, G., Jensen, K.C., Vogel, H., Xu, L.W., et al. (2014). Targeting a glioblastoma cancer stem-cell population defined by EGF receptor variant III. Cancer Res 74, 1238–1249.

14. Tchoghandjian, A., Baeza, N., Colin, C., Cayre, M., Metellus, P., Beclin, C., Ouafik, L., and Figarella-Branger, D. (2010). A2B5 cells from human glioblastoma have cancer stem cell properties. Brain Pathol 20, 211–221.

15. Patel, A.P., Tirosh, I., Trombetta, J.J., Shalek, A.K., Gillespie, S.M., Wakimoto, H., Cahill, D.P., Nahed, B.V., Curry, W.T., Martuza, R.L., et al. (2014). Single-cell RNA-seq highlights intratumoral heterogeneity in primary glioblastoma. Science 344, 1396–1401.

16. Bao, S., Wu, Q., McLendon, R.E., Hao, Y., Shi, Q., Hjelmeland, A.B., Dewhirst, M.W., Bigner, D.D., and Rich, J.N. (2006). Glioma stem cells promote radioresistance by preferential activation of the DNA damage response. Nature 444, 756–760.

17. Chen, J., Li, Y., Yu, T.S., McKay, R.M., Burns, D.K., Kernie, S.G., and Parada, L.F. (2012). A restricted cell population propagates glioblastoma growth after chemotherapy. Nature 488, 522–526.

18. Batista, P.J., and Chang, H.Y. (2013). Long noncoding RNAs: cellular address codes in development and disease. Cell 152, 1298–1307.

19. Beermann, J., Piccoli, M.T., Viereck, J., and Thum, T. (2016). Non-coding RNAs in Development and Disease: Background, Mechanisms, and Therapeutic Approaches. Physiol Rev 96, 1297–1325.

20. Slack, F.J., and Chinnaiyan, A.M. (2019). The Role of Non-coding RNAs in Oncology. Cell 179, 1033–1055.

21. Consortium, E.P., Birney, E., Stamatoyannopoulos, J.A., Dutta, A., Guigo, R., Gingeras, T.R., Margulies, E.H., Weng, Z., Snyder, M., Dermitzakis, E.T., et al. (2007). Identification and analysis of functional elements in 1% of the human genome by the ENCODE pilot project. Nature 447, 799–816.

22. Carninci, P., Kasukawa, T., Katayama, S., Gough, J., Frith, M.C., Maeda, N., Oyama, R., Ravasi, T., Lenhard, B., Wells, C., et al. (2005). The transcriptional landscape of the mammalian genome. Science 309, 1559–1563.

23. Katayama, S., Tomaru, Y., Kasukawa, T., Waki, K., Nakanishi, M., Nakamura, M., Nishida, H., Yap, C.C., Suzuki, M., Kawai, J., et al. (2005). Antisense transcription in the mammalian transcriptome. Science 309, 1564–1566.

24. Rinn, J.L., and Chang, H.Y. (2012). Genome regulation by long noncoding RNAs. Annu Rev Biochem 81, 145–166.

25. Wang, K.C., and Chang, H.Y. (2011). Molecular mechanisms of long noncoding RNAs. Mol Cell 43, 904–914.

26. Ulitsky, I., and Bartel, D.P. (2013). lincRNAs: genomics, evolution, and mechanisms. Cell 154, 26–46.

27. Guttman, M., Garber, M., Levin, J.Z., Donaghey, J., Robinson, J., Adiconis, X., Fan, L., Koziol, M.J., Gnirke, A., Nusbaum, C., et al. (2010). Ab initio reconstruction of cell type-specific transcriptomes in mouse reveals the conserved multi-exonic structure of lincRNAs. Nat Biotechnol 28, 503–510.

28. Guttman, M., Donaghey, J., Carey, B.W., Garber, M., Grenier, J.K., Munson, G., Young, G., Lucas, A.B., Ach, R., Bruhn, L., et al. (2011). lincRNAs act in the circuitry controlling pluripotency and differentiation. Nature 477, 295–300.

29. Yao, R.W., Wang, Y., and Chen, L.L. (2019). Cellular functions of long noncoding RNAs. Nat Cell Biol 21, 542–551.

30. Statello, L., Guo, C.J., Chen, L.L., and Huarte, M. (2021). Gene regulation by long non-coding RNAs and its biological functions. Nat Rev Mol Cell Biol 22, 96–118.

31. Bergmann, J.H., Li, J., Eckersley-Maslin, M.A., Rigo, F., Freier, S.M., and Spector, D.L. (2015). Regulation of the ESC transcriptome by nuclear long noncoding RNAs. Genome Res 25, 1336–1346.

32. Yang, G., Lu, X., and Yuan, L. (2014). LncRNA: a link between RNA and cancer. Biochim Biophys Acta 1839, 1097–1109.

33. Liu, S.J., Nowakowski, T.J., Pollen, A.A., Lui, J.H., Horlbeck, M.A., Attenello, F.J., He, D., Weissman, J.S., Kriegstein, A.R., Diaz, A.A., et al. (2016). Single-cell analysis of long non-coding RNAs in the developing human neocortex. Genome Biol 17, 67.

34. Curry, R.N., and Glasgow, S.M. (2021). The Role of Neurodevelopmental Pathways in Brain Tumors. Front Cell Dev Biol 9, 659055.

35. Han, Y., Zhou, L., Wu, T., Huang, Y., Cheng, Z., Li, X., Sun, T., Zhou, Y., and Du, Z. (2016). Downregulation of lncRNA-MALAT1 Affects Proliferation and the Expression of Stemness Markers in Glioma Stem Cell Line SHG139S. Cell Mol Neurobiol 36, 1097–1107.

36. Yao, Y., Ma, J., Xue, Y., Wang, P., Li, Z., Liu, J., Chen, L., Xi, Z., Teng, H., Wang, Z., et al. (2015). Knockdown of long non-coding RNA XIST exerts tumor-suppressive functions in human glioblastoma stem cells by up-regulating miR-152. Cancer Lett 359, 75–86.

37. Gong, W., Zheng, J., Liu, X., Ma, J., Liu, Y., and Xue, Y. (2016). Knockdown of NEAT1 restrained the malignant progression of glioma stem cells by activating microRNA let-7e. Oncotarget 7, 62208–62223.

38. Su, R., Cao, S., Ma, J., Liu, Y., Liu, X., Zheng, J., Chen, J., Liu, L., Cai, H., Li, Z., et al. (2017). Knockdown of SOX2OT inhibits the malignant biological behaviors of glioblastoma stem cells via up-regulating the expression of miR-194-5p and miR-122. Mol Cancer 16, 171.

39. Jiang, X., Yan, Y., Hu, M., Chen, X., Wang, Y., Dai, Y., Wu, D., Wang, Y., Zhuang, Z., and Xia, H. (2016). Increased level of H19 long noncoding RNA promotes invasion, angiogenesis, and stemness of glioblastoma cells. J Neurosurg 2016, 129–136.

40. Bhaduri, A., Di Lullo, E., Jung, D., Muller, S., Crouch, E.E., Espinosa, C.S., Ozawa, T., Alvarado, B., Spatazza, J., Cadwell, C.R., et al. (2020). Outer Radial Glia-like Cancer Stem Cells Contribute to Heterogeneity of Glioblastoma. Cell Stem Cell 26, 48–63 e46.

41. Jacob, F., Salinas, R.D., Zhang, D.Y., Nguyen, P.T.T., Schnoll, J.G., Wong, S.Z.H., Thokala, R., Sheikh, S., Saxena, D., Prokop, S., et al. (2020). A Patient-Derived Glioblastoma Organoid Model and Biobank Recapitulates Inter- and Intra-tumoral Heterogeneity. Cell 180, 188–204 e122.

42. Nowakowski, T.J., Bhaduri, A., Pollen, A.A., Alvarado, B., Mostajo-Radji, M.A., Di Lullo, E., Haeussler, M., Sandoval-Espinosa, C., Liu, S.J., Velmeshev, D., et al. (2017). Spatiotemporal gene expression trajectories reveal developmental hierarchies of the human cortex. Science 358, 1318–1323.

43. Wang, R., Sharma, R., Shen, X., Laughney, A.M., Funato, K., Clark, P.J., Shpokayte, M., Morgenstern, P., Navare, M., Xu, Y., et al. (2020). Adult Human Glioblastomas Harbor Radial Glia-like Cells. Stem Cell Reports 14, 338–350.

44. Zhao, S.G., Yu, M., Spratt, D.E., Chang, S.L., Feng, F.Y., Kim, M.M., Speers, C.W., Carlson, B.L., Mladek, A.C., Lawrence, T.S., et al. (2019). Xenograft-based, platform-independent gene signatures to predict response to alkylating chemotherapy, radiation, and combination therapy for glioblastoma. Neuro Oncol 21, 1141–1149.

45. Hazra, R., Brine, L., Garcia, L., Benz, B., Chirathivat, N., Shen, M.M., Wilkinson, J.E., Lyons, S.K., and Spector, D.L. (2022). Platr4 is an early embryonic lncRNA that exerts its function downstream on cardiogenic mesodermal lineage commitment. Dev Cell 57, 2450–2468 e2457.

46. Hazra, R., and Spector, D.L. (2022). Simultaneous visualization of RNA transcripts and proteins in whole-mount mouse preimplantation embryos using single-molecule fluorescence in situ hybridization and immunofluorescence microscopy. Front Cell Dev Biol 10, 986261.

47. Gimple, R.C., Bhargava, S., Dixit, D., and Rich, J.N. (2019). Glioblastoma stem cells: lessons from the tumor hierarchy in a lethal cancer. Genes Dev 33, 591–609.

48. Alvarado, A.G., Thiagarajan, P.S., Mulkearns-Hubert, E.E., Silver, D.J., Hale, J.S., Alban, T.J., Turaga, S.M., Jarrar, A., Reizes, O., Longworth, M.S., et al. (2017). Glioblastoma Cancer Stem Cells Evade Innate Immune Suppression of Self-Renewal through Reduced TLR4 Expression. Cell Stem Cell 20, 450–461 e454.

49. Vaubel, R.A., Tian, S., Remonde, D., Schroeder, M.A., Mladek, A.C., Kitange, G.J., Caron, A., Kollmeyer, T.M., Grove, R., Peng, S., et al. (2020). Genomic and Phenotypic Characterization of a Broad Panel of Patient-Derived Xenografts Reflects the Diversity of Glioblastoma. Clin Cancer Res 26, 1094–1104.

50. Andersen, R.E., and Lim, D.A. (2018). Forging our understanding of lncRNAs in the brain. Cell Tissue Res 371, 55–71.

51. Lendahl, U., Zimmerman, L.B., and McKay, R.D. (1990). CNS stem cells express a new class of intermediate filament protein. Cell 60, 585–595.

52. Novak, D., Huser, L., Elton, J.J., Umansky, V., Altevogt, P., and Utikal, J. (2020). SOX2 in development and cancer biology. Semin Cancer Biol 67, 74–82.

53. Holmberg Olausson, K., Maire, C.L., Haidar, S., Ling, J., Learner, E., Nister, M., and Ligon, K.L. (2014). Prominin-1 (CD133) defines both stem and non-stem cell populations in CNS development and gliomas. PLoS One 9, e106694.

54. Cheetham, S.W., Gruhl, F., Mattick, J.S., and Dinger, M.E. (2013). Long noncoding RNAs and the genetics of cancer. Br J Cancer 108, 2419–2425.

55. Huarte, M. (2015). The emerging role of lncRNAs in cancer. Nat Med 21, 1253–1261.

56. Diermeier, S.D., Chang, K.C., Freier, S.M., Song, J., El Demerdash, O., Krasnitz, A., Rigo, F., Bennett, C.F., and Spector, D.L. (2016). Mammary Tumor-Associated RNAs Impact Tumor Cell Proliferation, Invasion, and Migration. Cell Rep 17, 261–274.

57. Chen, Y., Hao, Q., Wang, S., Cao, M., Huang, Y., Weng, X., Wang, J., Zhang, Z., He, X., Lu, H., et al. (2021). Inactivation of the tumor suppressor p53 by long noncoding RNA RMRP. Proc Natl Acad Sci U S A 118.

58. Liu, Z., Wang, X., Yang, G., Zhong, C., Zhang, R., Ye, J., Zhong, Y., Hu, J., Ozal, B., and Zhao, S. (2020). Construction of lncRNA-associated ceRNA networks to identify prognostic lncRNA biomarkers for glioblastoma. J Cell Biochem 121, 3502–3515.

59. Liu, S., Mitra, R., Zhao, M.M., Fan, W., Eischen, C.M., Yin, F., and Zhao, Z. (2016). The Potential Roles of Long Noncoding RNAs (lncRNA) in Glioblastoma Development. Mol Cancer Ther 15, 2977–2986.

60. Paul, Y., Thomas, S., Patil, V., Kumar, N., Mondal, B., Hegde, A.S., Arivazhagan, A., Santosh, V., Mahalingam, K., and Somasundaram, K. (2018). Genetic landscape of long noncoding RNA (lncRNAs) in glioblastoma: identification of complex lncRNA regulatory networks and clinically relevant lncRNAs in glioblastoma. Oncotarget 9, 29548–29564.

61. Reon, B.J., Anaya, J., Zhang, Y., Mandell, J., Purow, B., Abounader, R., and Dutta, A. (2016). Expression of lncRNAs in Low-Grade Gliomas and Glioblastoma Multiforme: An In Silico Analysis. PLoS Med 13, e1002192.

62. He, Y., Wang, L., Tang, J., and Han, Z. (2020). Genome-Wide Identification and Analysis of the Methylation of lncRNAs and Prognostic Implications in the Glioma. Front Oncol 10, 607047.

63. Zhu, X., Chen, H.H., Gao, C.Y., Zhang, X.X., Jiang, J.X., Zhang, Y., Fang, J., Zhao, F., and Chen, Z.G. (2020). Energy metabolism in cancer stem cells. World J Stem Cells 12, 448–461.

